# The draft genome sequence of Japanese rhinoceros beetle *Trypoxylus dichotomus*

**DOI:** 10.1101/2022.01.10.475740

**Authors:** Shinichi Morita, Tomoko F. Shibata, Tomoaki Nishiyama, Yuuki Kobayashi, Katsushi Yamaguchi, Kouhei Toga, Takahiro Ohde, Hiroki Gotoh, Takaaki Kojima, Jesse Weber, Marco Salvemini, Takahiro Bino, Mutsuki Mase, Moe Nakata, Tomoko Mori, Shogo Mori, Richard Cornette, Kazuki Sakura, Laura C. Lavine, Douglas J. Emlen, Teruyuki Niimi, Shuji Shigenobu

## Abstract

Beetles are the largest insect order and one of the most successful animal groups in terms of number of species. The Japanese rhinoceros beetle *Trypoxylus dichotomus* (Coleoptera, Scarabaeidae, Dynastini) is a giant beetle with distinctive exaggerated horns present on the head and prothoracic regions of the male. *T. dichotomus* has been used as research model in various fields such as evolutionary developmental biology, ecology, ethology, biomimetics, and drug discovery. In this study, *de novo* assembly of 615 Mb, representing 80% of the genome estimated by flow cytometry, was obtained using the 10x Chromium platform. The scaffold N50 length of the genome assembly was 8.02 Mb, with repetitive elements predicted to comprise 49.5% of the assembly. In total, 23,987 protein-coding genes were predicted in the genome. In addition, *de novo* assembly of the mitochondrial genome yielded a contig of 20,217 bp. We also analyzed the transcriptome by generating 16 RNA-seq libraries from a variety of tissues of both sexes and developmental stages, which allowed us to identify 13 co-expressed gene modules. The detailed genomic and transcriptomic information of *T. dichotomus* is the most comprehensive among those reported for any species of Dynastinae. This genomic information will be an excellent resource for further functional and evolutionary analyses, including the evolutionary origin and genetic regulation of beetle horns and the molecular mechanisms underlying sexual dimorphism.

## Introduction

Beetles (Insecta: Coleoptera) are the largest order not only among insects but among all animal, and they are regarded as one of the most successful animal groups.^1^ Beetles exhibit extraordinary morphological, ecological, and behavioral diversity^2^ and have been used as models to study ecological and evolutionary biology for centuries. For example, when Darwin first introduced the term “sexual selection,” the rhinoceros beetle horn was featured as a typical example.^3^

*Trypoxylus dichotomus*, commonly known as the Japanese rhinoceros beetle, is a giant beetle reaching up to 91.7 mm in length (Fig. 1a). First described by Linnaeus,^4^ *T. dichotomus* belongs to the Scarabaeidae, Dynastini, Trypoxylus. The genus Trypoxylus is most closely related to the genus Xyloscaptes (Fig. 1b).^1,5–8^ *T. dichotomus* inhabits East Asia including Japan, China, Tibet, Taiwan, the Korean peninsula, Northeastern India, Thailand, Vietnam, Laos, and Myanmar. *T. dichotomus* is characterized by the set of exaggerated horns present in the head and prothoracic regions of males, while females have no horns. The horn of the head region (head horn) can extend to more than 2/3 of the male’s body length and is bifurcated twice at the distal tip. The horn of the prothoracic region (thoracic horn) is shorter and bifurcated once at the distal tip. Both horns are strongly sexually dimorphic, and are used as weapons in combat between males over feeding territories visited by females.^9^ Battles take place on the trunks of host trees, with males inserting their head horn under the prothorax of an opponent and attempting to scoop the rival off of the tree. Biomechanical studies of horn morphology indicate that the shape of the horn, particularly the triangular cross-section of the base of the horn, resists buckling when twisted and is well suited to the nature of the battles in this species.^10^

**Figure 1.**
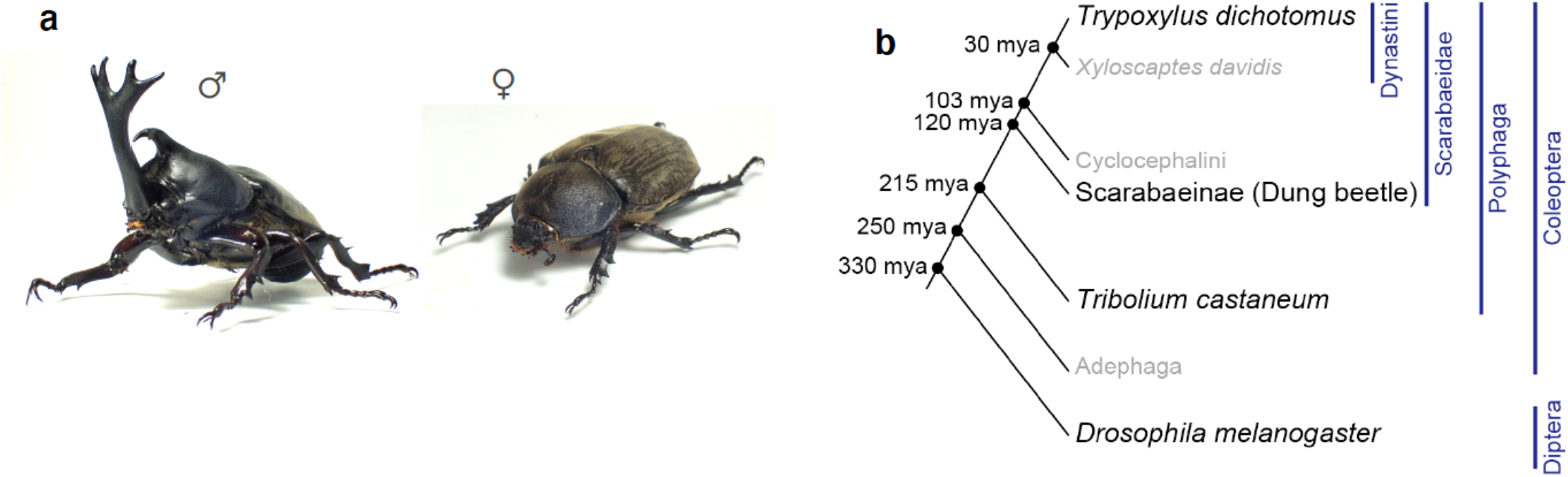
Photograph and phylogenetic context of the Japanese rhinoceros beetle, *Trypoxylus dichotomus*. (a) Photograph of an adult male (left) and an adult female (right) of *T. dichotomus*. (b) A phylogenetic tree of beetles depicting the phylogenetic relationship between *T. dichotomus* and related beetles with *Drosophila melanogaste*r as an outgroup. Estimated divergence dates (mya: million years ago) are based on Hunt et al., 2007, Ahrens et al., 2014, Misof et al., 2014, Mckenna et al., 2015 and Jin et al., 2016.^1,5–8^

*T. dichotomus* is a univoltine insect. Larvae feed and grow on humus soil or decaying wood from autumn to spring. During the prepupal stage, the larvae form a pupal chamber using a mixture of fecal pellets and humus to pupate below ground. In early summer, the adults appear in broad-leaved forests to ingest sap, and males and females mate on trees near feeding sites. After mating, the female lays the eggs in the humus. Adults have a maximum lifespan of about three months and are unable to overwinter.

*T. dichotomus* is an emerging model insect with many advantages. First, it is easy to obtain and the breeding / culturing system in the laboratory has been well developed. Since *T. dichotomus* is a popular pet in Japan, larvae are commercially available for approximately 1 USD each, and related items necessary for breeding (e.g., breeding cages, food for larvae and adults, artificial pupal chambers, etc.) can be purchased at a low cost. Larvae are easily maintained in the laboratory and can be stored at low temperatures to delay the onset of pupation, facilitating their use for research throughout the year.^11^ For the critical period from prepupa to adult, when horn formation occurs, a soil-free breeding system has been established that permits non-invasive continuous monitoring and the precise staging of horn development.^12^ Second, an RNA interference (RNAi) technique has been established in *T. dichotomus*, which allows functional assessment of genes of interest.^11–16^ Larval RNAi is performed by injecting double-stranded RNA (dsRNA) into the 1st thoracic segment (T1 segment) of the last instar larva just before the prepupal stage. Larval RNAi is so efficient that researchers can carry out large-scale RNAi screens against candidate genes, as proved by our successful identification of genes related to horn formation.^11–16^ Third, the large body size of *T. dichotomus* has advantages in efficient sampling for various experiments, which could facilitate biochemical assays and molecular analyses including next-generation sequencing (NGS).^15,17^ For example, sufficient amounts of RNA and DNA required for NGS studies can be obtained from each tissue of a single individual alone as shown in this paper.

Studies of *T. dichotomus* have increased rapidly in a wide variety of research fields due to the utility of this species as a model system. For example, in developmental biology the mechanisms of *T. dichotomus* horn formation have been described in depth.^18^ The molecular pathways underlying sexual dimorphism of the exaggerated horns have been intensively studied,^11,12^ and numerous genes involved in horn formation have been identified by comprehensive transcriptome analyses.^15^ In addition, beetle horns are considered to be an evolutionary novelty because there is no obvious homologous structure in the ancestral species,^19^ and *T. dichotomus* horns have proven to be an excellent model for exploring how novel morphological structures arise.

The length of *T. dichotomus* horns is exquisitely sensitive to the nutritional condition of larvae.^20^ Although many exaggerated ornaments and weapons of sexual selection exhibit such nutrition- or condition-dependent expression – indeed, this plasticity is considered to be an integral component of their function as a reliable signal of the quality of a male -- the horns of *T. dichotomus* were the first sexually-selected structure of any animal species to have these underlying mechanisms of conditional expression explored at a developmental or genetic level.^13^ The resulting variation in male horn length is mildly polyphenic, yielding major and minor males,^9,21^ and ethological studies revealed that males tap each other with their head horns before direct combat,^22^ to assess the size of their competitors and avoid unnecessary fighting.^23^ Less competitive minor males have alternative reproductive strategies to spend more time mating with females^24^ and appear at feeding sites earlier than major males to avoid fighting and encounter females.^25,26^

The large size and extraordinary morphology of *T. dichotomus* has inspired several biomimetic studies. For example, the mechanical properties of elytra (fore wings)^27–30^ and the aerodynamic mechanisms^31^ of *T. dichotomus* flight have been studied and applied to industrial uses. In addition, an anti-bacterial peptide^32–38^ and a molecule with anti-prion activity^39^ were discovered from *T. dichotomus* and medical applications of these molecules are expected. Thus, *T. dichotomus* has been actively used in various research fields, including evolutionary developmental biology, ecology, ethology, biomimetics, and drug discovery.

Despite the excellent properties of this beetle as a research model, limited genetic information is available for *T. dichotomus*, although several NGS analyses were reported.^15,17,40^ In the present study, we present the draft genome of *T. dichotomus*, together with the functional annotation and gene expression data. In addition, we compared the gene repertoire of *T. dichotomus* with those of three other insects. This genomic information provides a crucial foundation for future studies using *T. dichotomus* as well as the comparative genomics of insects.

## Materials and Methods

### Insects

We purchased *T. dichotomus* larvae from Loiinne (Gunma, Japan). Last (third) instar larvae were sexed as described previously,^11^ individually fed on humus in plastic containers, and kept at 10°C until use. For tissue sampling, larvae were moved to room temperature for a minimum of 10 days and reared at 28 °C.

### Genome sequencing and *de novo* assembly

Male leg primordia derived from a single individual at 72 h after pupal-chamber formation, which is before cuticle pigmentation and sclerotization, were dissected out from prepupa in 0.75% sodium chloride solution, and used for genome analysis.

To prepare high molecular weight (HMW) DNA of the beetle, 184.5 mg of the frozen tissue was transferred to a mortar and gently ground into a fine powder with liquid nitrogen. Frozen powdery QIAGEN G2 buffer (Qiagen, Netherlands, Cat #1014636), which was generated by spraying the buffer into liquid nitrogen in a glass beaker, was added to the sample and blended quickly. Letting the mixture thaw in a tube, RNaseA (QIAGEN, Cat #19101) and Proteinase K (QIAGEN, Cat #1019499) were added, and the sample was incubated at room temperature for 2.5 h without agitation. The sample was centrifuged at 5000 x g for 30 min and the supernatant was subjected to DNA extraction with a QIAGEN Genomic-tip 100/G column (Cat # 10243). The genomic DNA was eluted with 5 ml of Buffer QF, and then purified and concentrated using 0.5-fold volume of Agencourt AMPure XP magnetic beads (Beckman Coulter, Brea, CA, USA, Cat # A63881) yielding 12.2 µg of DNA (174.0 ng/µl). The DNA was size-fractioned by using SAGE HLS (Sage Science, Beverly, MA, USA, Cat #HLS0001), and the fraction with the highest molecular size was collected. The size distribution of the HMW DNA was evaluated by pulsed-field-gel-electrophoresis (PFGE). In brief, 20 ng of HMW DNA was run on a 1% agarose gel (Seakem Gold Agarose, Lonza, Rockland, ME, USA, Cat #50150) in 0.5xTBE with the BioRad CHEF Mapper system (BioRad, Hercules, CA, USA, Cat #M1703650) for 15 h, and the gel was stained with SYBR Gold dye (Thermo Fisher Scientific, Waltham, MA, USA, Cat #11494). Lambda ladder, *Saccharomyces cerevisiae* genome, and 5 kbp ladder (BioRad,Cat #170–3624) were used as standards. These results demonstrated the HMW DNA had an approximate mean size of 50 - 80 kbp, while shorter DNA fragments (< ∼10 kbp) were efficiently removed. DNA concentration was measured with a Qubit fluorometer (Thermo Fisher Scientific, Cat #Q32866) using the Qubit(tm) dsDNA BR Assay Kit (Thermo Fisher Scientific, Cat #Q32850).

We constructed a 10x Genomics Chromium linked-read library using 0.68 ng of the HMW DNA extracted as described above with a Chromium Genome Library Kit & Gel Bead Kit v2 (10x Genomics, San Francisco, CA, USA, Cat #120258) following the manufacturer’s protocol. The generated library was sequenced on a HiSeqX (Ilumina, San Diego, CA, USA) at Macrogen Japan Corp (Tokyo, Japan). The obtained Illumina reads were assembled using the Supernova assembler (ver. 2.1.0)^41^ with a parameter of maxreads = 265000000.

We assembled the mitochondrial genome of *T. dichotomus* separately from a paired-end library. Genomic DNA was extracted from the leg primordia using the QIAGEN Genomic-Tip. A paired-end library was prepared with the TruSeq DNA PCR-Free Library Preparation Kit (Illumina) from 1 µg of the genomic DNA. The library was sequenced using the Illumina HiSeqX system with 2 × 150 bp paired-end sequencing protocol at Macrogen Japan Corp (Tokyo, Japan), where 182,102,568 read-pairs were produced. The mitochondrial genome was assembled using a subset of the raw reads (1 M reads, 302 Mb) by NovoPlasty (v4.3.1)^42^ with a parameter of K-mer = 33, using a partial sequence of 16S rRNA of *T. dichotomus* (GenBank accession: AB178318.1) as the seed sequence. Our *de novo* assembly yielded two contigs, Contig1 (20,217 bp) and Contig2 (1,163 bp). A manual inspection revealed that Contig2 was a repetitive sequence included in Contig1 and Contig1 corresponds to the mitochondrial genome. Note that the repetitive regions were not fully resolved by the *de novo* assembly from Illumina short reads and it is likely the contig, in fact, contains more units of the repeats. The assembled contig was annotated using MITOS2 webserver (http://mitos2.bioinf.uni-leipzig.de)^43^ with the invertebrate mitochondrial genetic code.

### Genome annotation

Repeat annotation: We analyzed the distribution of repetitive elements in the genome of *T. dichotomus*, using RepeatModeler (ver. open-1.0.8)^44^ and RepeatMasker (ver. open-4.0.6)^45^.

Gene prediction: We annotated the *T. dichotomu*s genome for protein-coding genes using RNA-seq data. We generated 16 libraries covering 6 tissues and 6 developmental time points from early embryogenesis to postembryonic stages (see *RNA-Seq analysis* section below). The Illumina RNA-seq reads were aligned against a hard-masked genome in which genomic repetitive elements were substituted by ‘N’, using HISAT2 (ver. 2.1.0)^46^ with default parameters. The BRAKER2 (ver. 2.1.5)^47^ pipeline was used to predict protein-coding genes based on the RNA-seq alignments.

Gene annotation: All predicted protein-coding genes were compared with the NCBI non-redundant protein database (nr DB, release November 2020) using the blastp command of diamond software (ver. 2.0.5)^48^ with a threshold of e-value < 1.0e-5. We used InterProScan (ver. 5.48)^49^ to query the predicted coding regions for known functional domains and assign Gene Ontology (GO) terms to the proteins. We also used the eggNOG-mapper pipeline (ver. 2.0.1b)^50^ for functional annotation (including GO term assignments) of the predicted genes based on the orthology information.

BUSCO (ver. 4.0.6)^51^ was used in quantitative measuring for the assessment of genome assembly, using insecta_odb10 (1367 total orthogroups) as the lineage input. A genome browser was built using JBrowse2 web (ver. 1.5.1)^52^ and is available at http://www.insect.nibb.info/trydi/.

### Ortholog analysis

We used the OrthoFinder (ver. 2.3.11)^53^ to generate clusters of orthologous and paralogous gene families. Public gene datasets of *Onthophagus taurus* (RefSeq accession No. GCF_000648695), *Tribolium castaneum* (UniProt accession No. UP000007266), and *Drosophila melanogaster* (UniProt accession No. UP000000803) were used as references.

### RNA-Seq analysis

In total, 16 samples (egg, non-sexed embryos from five stages, ovary, testis, and Malpighian tubule, hindgut, brain and fat body of males and females respectively (Table S1)) were used for RNA-seq analysis. Eggs, ovaries, and testes were dissected out from adult insects in 0.75% sodium chloride solution. Malpighian tubule, hindgut, brain and fat body were dissected out from third instar larvae in 0.75% sodium chloride solution. These samples were frozen in liquid nitrogen and stored at −80 °C until use.

Total RNA was extracted from each tissue sample, except for the fat body sample, using the RNeasy Micro Kit (QIAGEN, Cat # 74004) according to the manufacturer’s instructions. Total RNA of the fat bodies was extracted using TRIzol Reagent (Thermo Fisher Scientific, Cat #15596-026) according to the manufacturer’s instructions. These total RNAs were treated with DNase using RNase-Free DNase Set (QIAGEN, Cat #79254), from which Illumina sequencing libraries were prepared using the TruSeq Stranded mRNA library prep kit (llumina) following the manufacturer’s instructions. The libraries were multiplexed and sequenced using the Illumina HiSeq1500 system with 2 × 101 bp paired-end sequencing protocol at the Functional Genomics Facility of National Institute for Basic Biology.

*T. dichotomus* RNA-Seq reads generated from these 16 libraries were adapter-trimmed using Trim Galore! (ver. 0.5.0)^54^ and cutadapt (ver. 1.18)^55^. The cleaned RNA-Seq reads were mapped to the *T. dichotomus* genome assembly using HISAT2 (ver. 2.1.0)^46^ with default parameters. For quantification of the gene expression, StringTie (ver. 2.1.3)^56^ and prepDE.py (http://ccb.jhu.edu/software/stringtie/dl/prepDE.py) with a GTF file of the BRAKER2 gene prediction were used to generate the count matrix. The count data were normalized by the trimmed mean M values (TMM) method^57^ available in the edgeR library (ver. 3.32.1)^58^. To visualize profiles of gene expression, a multidimensional scaling (MDS) plot was generated using the edgeR software package.

To detect modules of co-expressed genes from the transcriptome data, Weighted Gene Correlation Network Analysis (WGCNA) was applied.^59^ Normalized count data were used for this analysis implemented in the WGCNA library (version 1.70)^60^. Genes expressed at low levels, and genes with low expression variance across the libraries, were filtered out; the 6666 surviving gene models were used in the WGCNA analysis. A signed network was constructed in WGCNA with specific parameter settings of power = 8, networkType=“signed”, TOMType=“unsigned”, minModuleSize=30.

### Genome size estimation using flow cytometry

Flow cytometry estimates were made by quantification of fluorescence from propidium iodide (PI) stained nuclei extracted from primordia of horns, legs or wings of male pupae with the fruitfly, *Drosophila melanogaster* (1C = 173.3 Mbp) as an internal standard. Samples were added to 1 mL of PBS and homogenized with BioMasher II (nippi, Tokyo, Japan, cat# 320102), and then 1 µl of 10% Triton-X (SIGMA-ALDRICH, St Louis, MO, USA, cat# 93443) and 4 µl of 100 mg/ml RNase A (QIAGEN, cat# 19101) were added to the homogenate. The resultant solution was passed through a 30 µm CellTrics filter (Sysmex Partec, Görlitz, Germany, cat# BP486257) and stained with 10 µg/ml PI (Sony Biotechnology, San Jose, CA, USA, cat# 2706505). The nuclei were analyzed on a Cell Sorter SH800 (SONY Biotechnology) for three replicates.

### Genome size estimation from reads of shotgun sequencing

Genomic DNA was extracted from the leg primordia using the QIAGEN Genomic-Tip (Cat # 10243). A paired-end library was prepared with the TruSeq DNA PCR-Free LT Sample Preparation Kit (Illumina, #FC-121-3001) from 1 µg of the genomic DNA. The library was sequenced using the Illumina HiSeqX system with 2 × 150 bp paired-end sequencing protocol at Macrogen Japan Corp (Tokyo, Japan), where 182,102,568 read-pairs were produced. To estimate the genome size of *T. dichotomus*, we analyzed the distribution of k-mers from the Illumina reads using Jellyfish (ver. 2.2.10)^61^ and GenomeScope 2.0 (git commit id: fdeb891)^62^. The distribution of kmers of size was analyzed with three different kmer sizes, 21, 31 and 41.

### Data availability

Data from whole-genome sequencing and transcriptome sequencing have been deposited in the DDBJ database under BioProject accession PRJDB12657. The analyzed data including genome assembly, gene prediction, annotation, and gene expression are available through FigShare (https://doi.org/10.6084/m9.figshare.c.5737754). The genome browser is available at http://www.insect.nibb.info/trydi/.

## Results and Discussion

### Genome size estimate

Using flow cytometry and the fruitfly *Drosophila melanogaster* as reference, we determined that the haploid genome size of *T. dichotomus* is 773.1 ± 24.6 Mb. In addition, we estimated the *T. dichotomus* genome size from the distribution of k-mers from Illumina reads of shotgun sequences. The distribution of kmers of size 21, 31 and 41 resulted in an estimated haploid genome size of 710–764 Mb. The small discrepancy between k-mer and cytometry-based estimates may be caused by the repetitive elements (see below), which can affect k-mer estimates.

### Genome assembly and evaluation

Genome sequencing of *T. dichotomus* was performed with genomic DNA isolated from leg primordia of a single male prepupa collected in Gunma, Japan. We prepared high molecular weight (HMW) DNA from the beetle sample with an approximate mean size of 50 - 80 kbp. The HMW genome DNA (0.68 ng) was subjected to the linked-read whole-genome sequencing library construction on the 10x Chromium platform. A draft genome was assembled with Supernova^41^ using 265.02 M Illumina reads (57.0 x coverage). The haploid genome assembly was named TdicSN1.0 and used for downstream analysis.

The final assembly TdicSn1.0 consists of 15,609 scaffolds with an N50 of 8.02 Mb and a total size of 615 Mb [Table 1], covering 80% of the genome. We evaluated the completeness of the assembly by using the benchmarking universal single-copy orthologs (BUSCO).^63^ The BUSCO analysis showed that our *T. dichotomus* genome assembly has high coverage of coding regions, capturing 99.4 % (98.9% complete, 0.5% fragmented) from the Insecta dataset (version 4.0.6; n = 1,367) [Table 2]. The score is comparable to the genome of the model beetle *Tribolium castaneum*.^64^

**Table 1.**
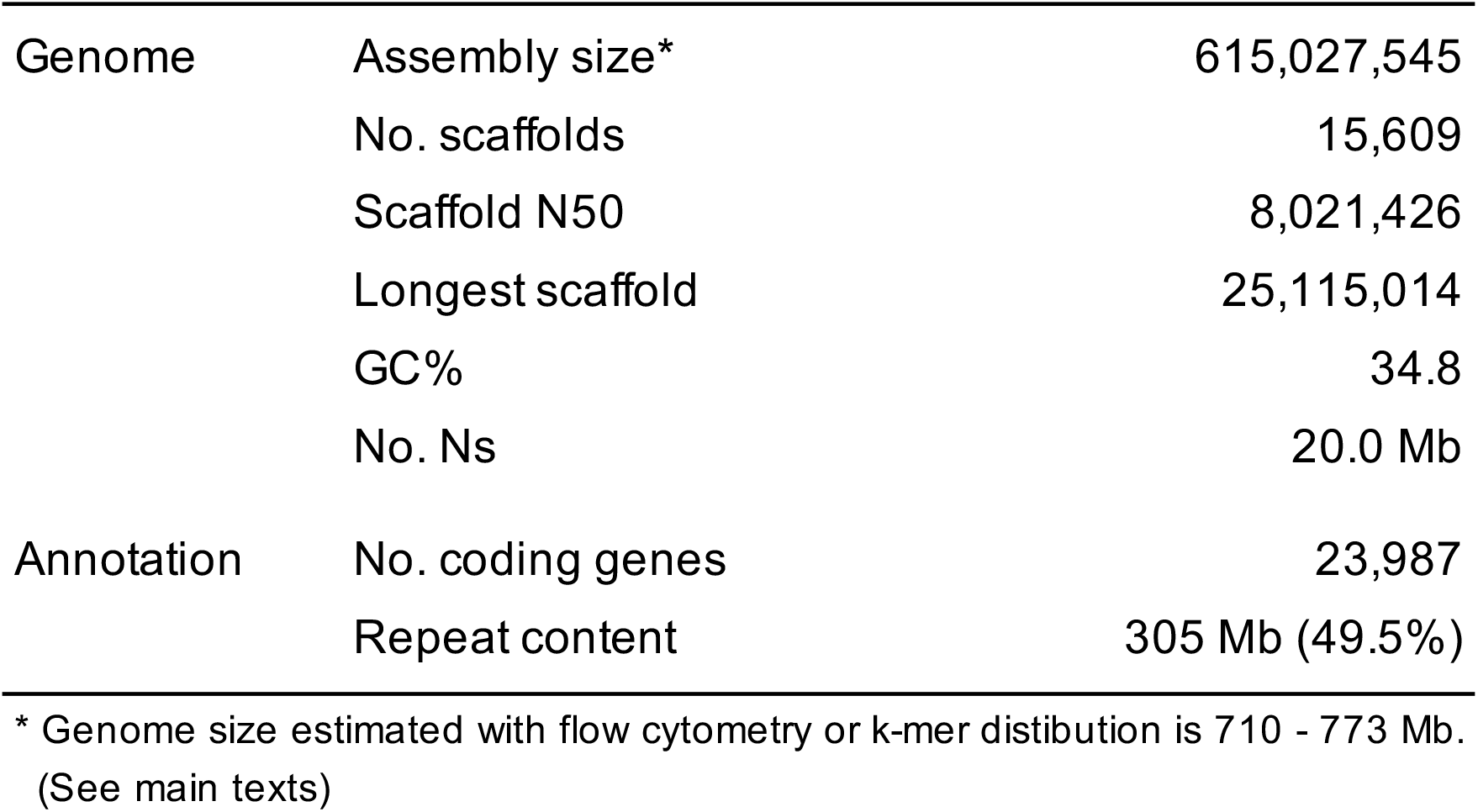
Summary of *T. dichotomus* genome assembly

**Table 2.** BUSCO analysis of the genome assembly of *T. dichotomus* using Insecta gene set.

| Database | insecta_odb10 |  |
| --- | --- | --- |
| Complete BUSCOs | 1352 | 98.9% |
| Complete and single-copy BUSCOs | 1335 | 97.7% |
| Complete and duplicated BUSCOs | 17 | 1.2% |
| Fragmented BUSCOs | 7 | 0.5% |
| Missing BUSCOs | 8 | 0.6% |
| Total BUSCO groups searched | 1367 |  |

We assembled the mitochondrial genome separately from a paired-end library using a known partial sequence of 16S rRNA of *T. dichotomus* (GenBank accession: AB178318.1) as the seed sequence. Our *de novo* assembly yielded a contig of 20,217 bp. We identified 2 ribosomal RNAs, 22 tRNAs, and 13 protein-coding genes. The gene repertoire and the structural arrangement showed typical features of mitochondrial genomes of insects, but the total length of the mitochondrial genome was longer than that of the model coleopteran *Tribolium castaneum* (15.8 kb (NC_003081.2)) by 4.3 kb. The increased length is mostly due to species specific repetitive elements between the s-rRNA gene and the tRNA-GLN gene.

### Genome annotation

The *T. dichotomus* genome is rich in repetitive elements, which total 305 Mb and account for 49.5% of the genome assembly [Table S2]. Considering the situation that the current assembly covers 80% of the estimated genome size, it is likely that uncovered regions contain more repetitive sequences that are generally difficult to capture by Illumina-based genome assembling. We annotated the *T. dichotomu*s genome for protein-coding genes using RNA-seq data. Aiming for a comprehensive gene identification, we collected RNA samples from a wide variety of tissues of both sexes and developmental stages. We generated 16 libraries derived from 6 tissues (brain, Malpighian tubule, hindgut, fat body, testis and ovary) and from 6 time points of whole embryos from middle to late developmental stages [Table S1]. The alignment data of these RNA-seq sequences mapped on the TdicSN1.0 assembly were subjected to the BRAKER2 pipeline to predict protein-coding genes. In total, 23,987 protein-coding genes are predicted in the *T. dichotomus* genome. Of these, 19,708 (82.6%) encoded proteins exhibiting sequence similarity to proteins in the NCBI non-redundant protein database. The two most frequent top-hit species corresponded to scarab beetles, *Oryctes borbonicus*, (6,312) and *Onthophagus taurus* (5,126), followed by other Coleoptera species such as *Tribolium castaneum, Ignelater luminosus*, and *Nicrophorus vespilloides*, which reasonably reflects the phylogenetic position of *T. dichotomus* [Fig 1b]. We used InterProScan to query the predicted coding regions for known functional domains. We identified 29,549 Pfam motifs^65^ and 34,594 PANTHER motifs^66^ in the products of 14,497 and 15,139 *T. dichotomus* gene models, respectively. We also identified 2,389 proteins with secretion potential predicted by the SignalP program^67^. By integrating protein-domain based and orthology-based approaches using InterProScan and eggNOG-mapper pipelines, 581,644 Gene Ontology terms were assigned to 10,541 genes (43.9%).

### Ortholog analysis

To understand the gene repertoire evolution of *T. dichotomus*, we generated clusters of orthologous and paralogous gene families comparing the *T. dichotomu*s proteome with those of three other insects, *Onthophagus taurus, Tribolium castaneum* and *Drosophila melanogaster. O. taurus* is a dung beetle belonging to the family Scarabaeidae [Fig. 1b], and represents another model insect for the study of horn polyphenism.^68^ We included *T. castaneum* as a model coleoptera and *D. melanogaster* as a model insect for comparison as outgroups. The OrthoFinder program identified 12,034 orthogroups consisting of 56,656 genes derived from all four insects. The 18,783 *T. dichotomus* gene models were clustered into 10,291 orthogroups, leaving 5,204 genes unassigned to any orthogroups (i.e., orphan genes). Among them, 8,219 groups were shared within three Coleoptera species, while 543 groups were shared within only the horned beetles, *O. taurus* and *T. dichotomus* [Fig. 2a]. These beetle-specific groups may account for the common characteristic traits such as the elytron of the beetle^69^ and the exaggerated horns of *T. dichotomus* and *O. taurus*. We found 967 groups, consisting of 5,705 genes, that are unique to *T. dichotomus* and may account for lineage specific traits [Fig. 2a].

**Figure 2.**
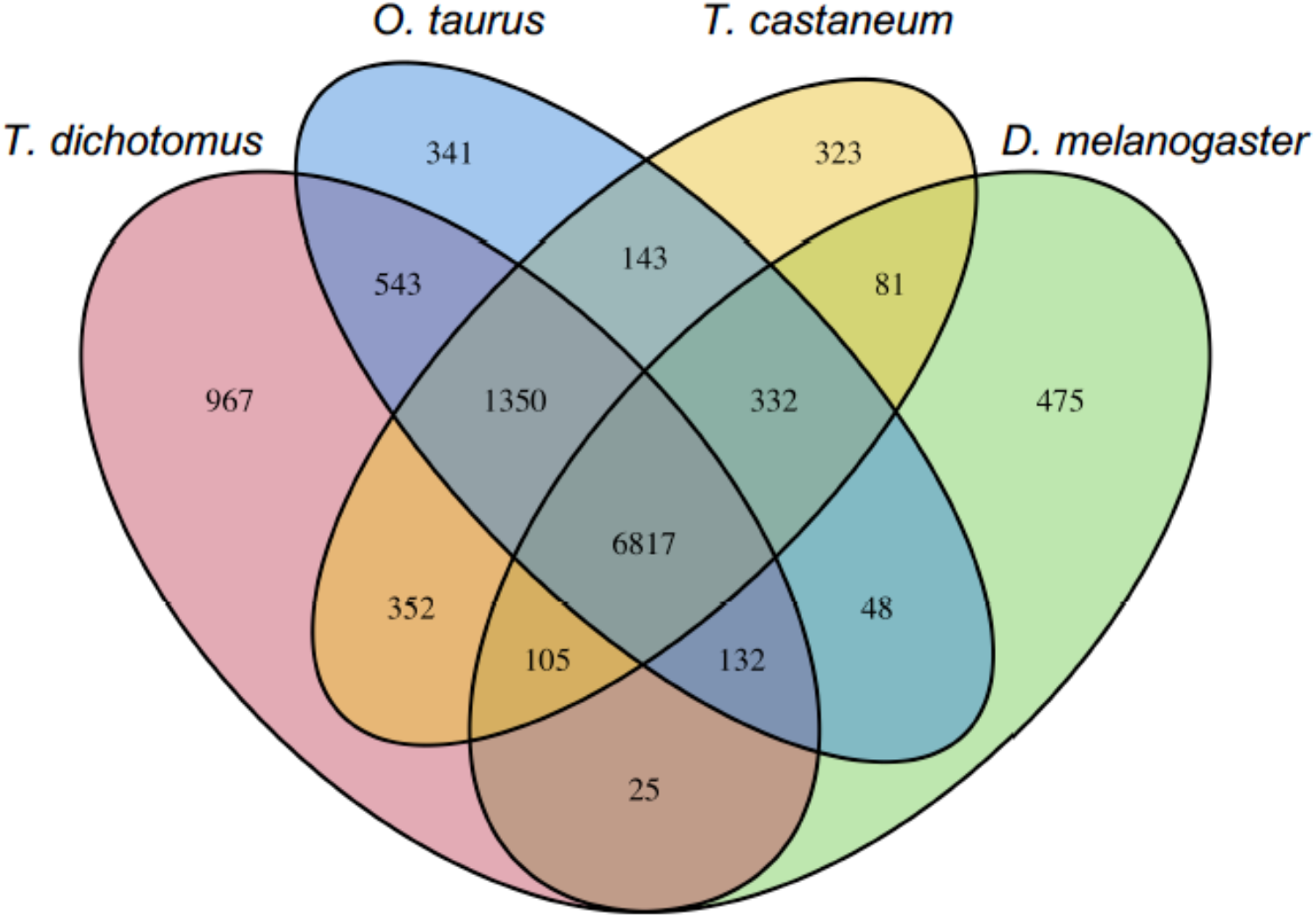
Venn diagram of shared and unique orthogroups in four insects. Orthogroups were identified by clustering of orthologous groups using OrthoFinder.^53^

### Transcriptome analysis

We analyzed 16 RNA-seq libraries created from a variety of tissues of both sexes and developmental stages [Table S1]. We quantified gene expression and profiled the expression patterns across all of the sample tissues. The multidimensional scaling (MDS) plot of the 16 samples depict the transcriptome similarities among the samples [Fig. 3a]. Samples derived from the same organs clustered together irrespective of sex differences. Eggs and ovaries also clustered together, as did the transcriptomes of embryos. Notably, embryonic samples were roughly ordered on the plot from earlier to later stages, which may represent the gradual transcriptional progression during embryogenesis of *T. dichotomus*.

**Figure 3.**
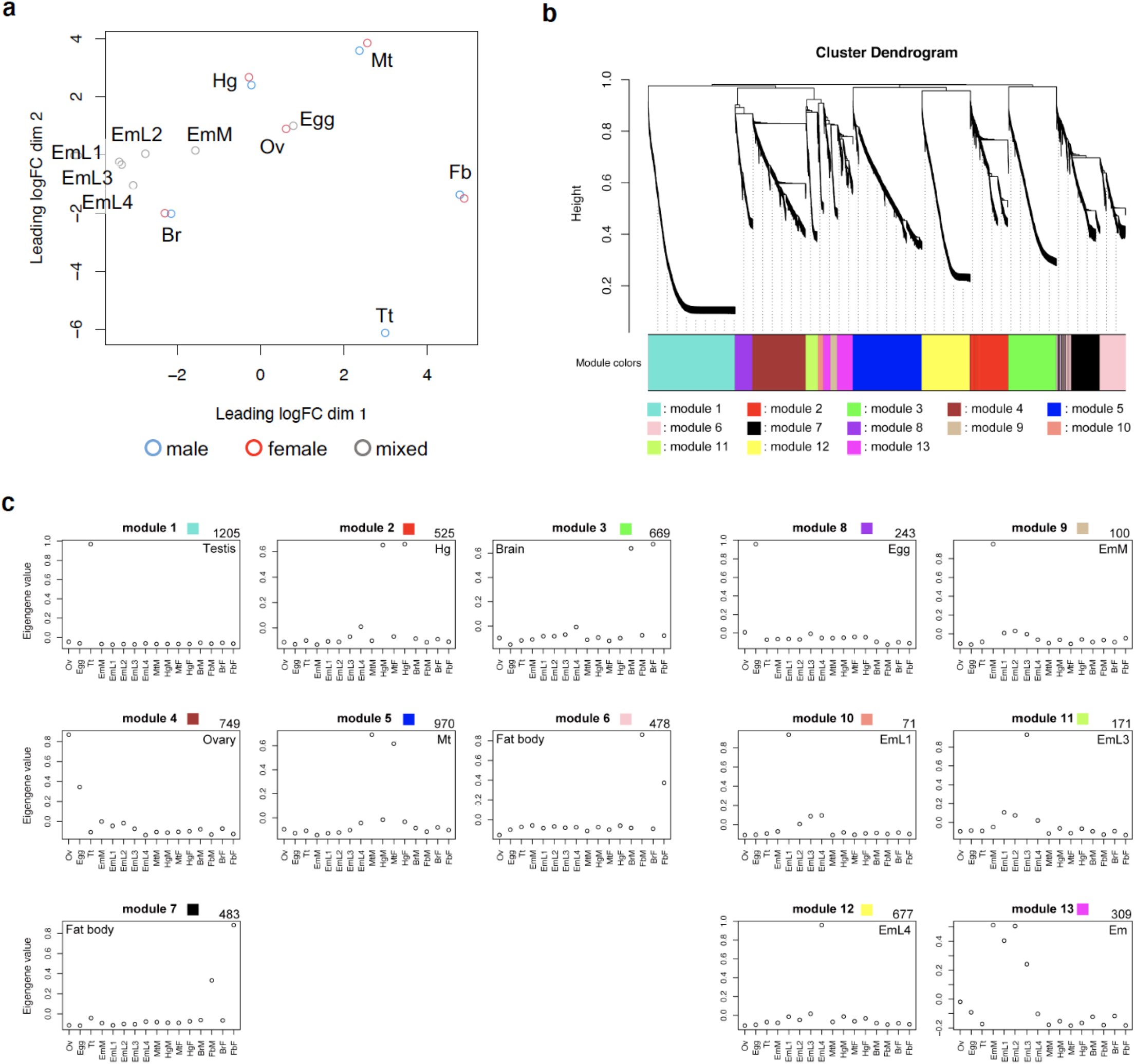
Transcriptome analysis. (a) MDS plot for RNA-Seq gene expression of *T. dichotomus* tissues, organs and developmental samples. Multi-dimensional scaling (MDS) plot showing relatedness between transcript expression profiles of 16 RNA-Seq libraries. Blue circles represent the expression profiles of male samples, and red circles represent those of female samples, while grey circles represent those of samples whose sex are unknown (i.e., embryos). The labels indicating the tissues and sources are defined as follows: Egg, eggs dissected out from the mature ovary; Tt, testis; Ov, ovary; EmM, middle-stage embryos at the stage of ventral appendage formation; EmL1, late stage embryo at the stage of appendages extend; EmL2, late-stage embryos at the stage of slightly more developed than EmL1; EmL3, late-embryos at a stage of the tracheal pits can be clearly observed; EmL4, late-embryos at a stage of the full-grown embryo; Mt, Malpighian tubules of third instar larva; Hg, hindgut tube of third instar larva; Fb, fat bodies of third instar larva; Br, brains of third instar larvae. See Table S1 for details of the label description. (b) Gene co-expression analysis of the *T. dichotomus* transcriptome. Hierarchical cluster tree of the *T. dichotomus* genes showing co-expression modules identified using WGCNA. Modules correspond to branches are labelled by colors as indicated by the color band underneath the tree. In total, 13 co-expression modules were identified from the expression data of 16 samples. (c) Co-expression gene modules identified using WGCNA are shown. Each circle represents the value of the respective module’s Eigengene. Module names shown above each panel correspond to (b). The sample abbreviations indicated by labels at the bottom of each panel are defined in the legend of (a). The number at the top right in each panel indicates the number of genes belonging to the module specified. The name of organ or developmental stage in which module genes preferentially expressed is displayed at top-right or top-left corner in each panel.

We performed Weighted Gene Correlation Network Analysis (WGCNA) to understand the co-expression relationship between genes at a system level. WGCNA identified 13 co-expressed modules from the expression data spanning 16 samples; each module contained 71 to 1205 co-expressed genes [Fig. 3bc]. Each module represents genes with highly correlated expression profiles, either in a single organ or in a certain stage of embryogenesis. Out of 13 modules, seven represent organ-specific patterns [Fig. 3bc, module 1 – 7]. Both modules 6 and 7 represent fat body preferential expression, but genes of the module 6 exhibited male-biased expression while genes in the module 7 exhibited female-biased expression, suggesting sexually differentiated functions of *T. dichotomus* fat body in the sexes. Six modules represent embryonic expression patterns [Fig. 3bc, module 8 – 13]. Genes contained in the module 13 exhibited a constant pattern of expression from middle to late stages.

## Conclusions

We report the assembly and annotation of the nuclear and mitochondrial genomes of the Japanese rhinoceros beetle *T. dichotomus*. The high completeness and the extensive annotations achieved for this genome, as well as transcriptome data generated from a wide variety of tissues and developmental stages, make this draft assembly an excellent resource for further functional and evolutionary analyses in this emerging model insect For example, the availability of the genome sequence will accelerate the use of functional assays using RNAi and genome editing techniques. These are key tools for experimental investigations of the evolutionary origin and genetic regulation of the exaggerated male horn, and the molecular mechanisms underlying the strong sexual dimorphism in its expression. We also anticipate that recent advances in genome sequencing such as long read sequencing and Hi-C proximity ligation methods will allow us extend the existing annotation to include a chromosome-level assembly of *T. dichotomus* in the near future, which would provide us with a more complete picture of the diploid genome composed of 18 autosomes and sex chromosomes (XX/XY).^70^ Other research areas such as ethology, biomimetics and drug discovery will benefit from the continuing advancement of genomic resources. Finally, given *T. dichotomus* is one of the most popular insects in Japan, we hope that the genomic resources will facilitate the development of effective population genomic tools to monitor and protect wild populations, which would lead to the establishment of the conservation genomics of the Japanese rhinoceros beetle. These genomic data and a genome browser are available at http://www.insect.nibb.info/trydi/.

## Acknowledgements

We thank Toshiya Ando, Taro Nakamura and Yasuhiko Chikami for helpful discussion, the Model Plant Research Facility/NIBB BioResource Center, the Data Integration and Analysis Facility/NIBB and the Functional Genomics Facility/NIBB Core Research Facilities for technical assistance. This work was supported by MEXT KAKENHI Grant Numbers 23128505, 25128706, 16H01452, 18H04766, 20H04933 and 20H05944 (to Te. N.); 17H06384 (to S.S.); 22128008 (to S.S. and To.N); JSPS KAKENHI Grant Numbers 19K16181 and 21K15135 (to S. M.), and NSF IOS–1456133 (to D. E. and L. L.).

## Tables

**Table S1.**
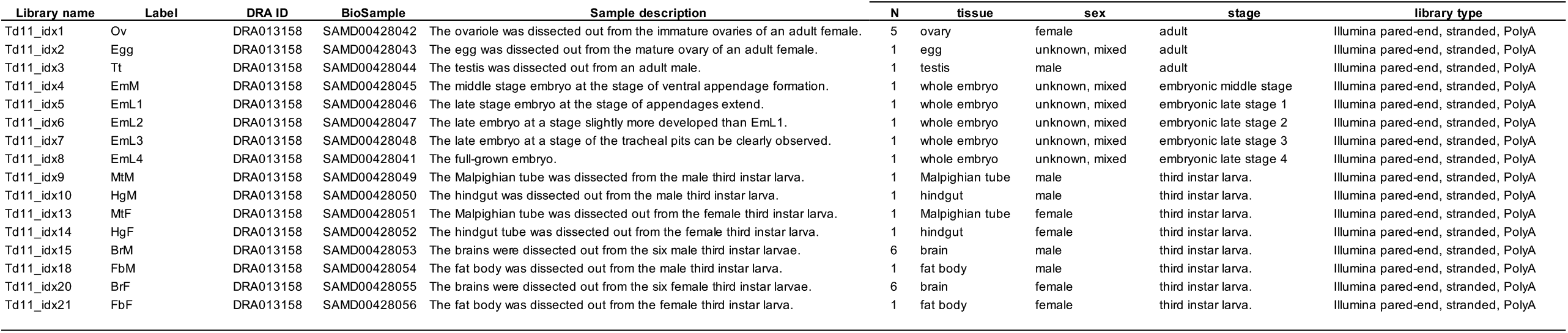
Samples used for RNA-seq

**Table S2.** Repetitive elements in the *T. dichotomus* genome.

|  | number of elements* | Total length (bp) | Percentage (%) |
| --- | --- | --- | --- |
| SINEs | 6798 | 758637 | 0.12 |
| ALUs | 0 | 0 | 0 |
| MIRs | 2487 | 273122 | 0.04 |
| LINEs | 137006 | 48445463 | 7.88 |
| LINE1 | 0 | 0 | 0 |
| LINE2 | 13502 | 4608407 | 0.75 |
| L3/CR1 | 36384 | 15348005 | 2.5 |
| LTR elements | 22213 | 5196007 | 0.84 |
| ERVL | 0 | 0 | 0 |
| ERVL-MaLRs | 0 | 0 | 0 |
| ERV_classI | 0 | 0 | 0 |
| ERV_classII | 0 | 0 | 0 |
| DNA elements | 503264 | 176254643 | 28.66 |
| hAT-Charlie | 10248 | 2803595 | 0.46 |
| TcMar-Tigger | 6485 | 2377631 | 0.39 |
| Unclassified: | 337209 | 70318994 | 11.43 |
| Total interspersed repeats |  | 300973744 | 48.94 |
| Small RNA: | 1229 | 194443 | 0.03 |
| Satellites: | 0 | 0 | 0 |
| Simple repeats: | 69943 | 3380637 | 0.55 |
| Low complexity: | 10747 | 543899 | 0.09 |
| Total repeat bases |  | 304697749 | 49.54 |

\* most repeats fragmented by insertions or deletions have been counted as one element

Data sets at FigShare (https://doi.org/10.6084/m9.figshare.c.5737754)

- Genome assembly: doi: 10.6084/m9.figshare.17126303
- Gene model: doi: 10.6084/m9.figshare.17126306
- Gene model annotation: 10.6084/m9.figshare.17126312
- RNA-seq data: 10.6084/m9.figshare.17126315
- RNA-seq WGCNA analysis: doi: 10.6084/m9.figshare.17126327

## Notes

### Competing Interest Statement

The authors have declared no competing interest.

